# Population structure and pangenome analysis of Enterobacter bugandensis uncover the presence of *bla*_CTX-M-55_, *bla*_NDM-5_ and *bla*_IMI-1_, along with sophisticated iron acquisition strategies

**DOI:** 10.1101/620682

**Authors:** Filipe P. Matteoli, Hemanoel Passarelli-Araujo, Francisnei Pedrosa-Silva, Fabio L. Olivares, Thiago M. Venancio

## Abstract

*Enterobacter bugandensis* is a recently described species that has been largely associated with nosocomial infections. Here, we report the genome of a non-clinical *E. bugandensis* strain. We used this and other several publicly available *E. bugandensis* genomes to obtain the species pangenome, investigate the conservation of important genes, and elucidate general population structure features of the species. Core- and whole-genome multilocus sequence typing (cgMLST and wgMLST, respectively) allowed the detection of five *E. bugandensis* phylogroups (PG-A to E). We found important antimicrobial resistance and virulence determinants associated with specific PGs, notably PG-A and PG-E. IncFII was the most prevalent plasmid replicon type in this species. We uncovered several extended-spectrum β-lactamases, including *bla*_CTX-M-55_ and *bla*_NDM-5_, present in an IncX replicon type plasmid, described here for the first time in *E. bugandensis*. Genetic context analysis of *bla*_NDM-5_ revealed the resemblance of this plasmid with other IncX plasmids isolated from other bacteria from the same country. Further, three distinctive siderophore producing operons were found in the *E. bugandensis* pangenome: enterobactin (*ent*), aerobactin (*iuc*/*iut*), and salmochelin (*iro*). The latter operon is conserved in all PG-E isolates. Collectively, our findings provide novel insights on the lifestyle, physiology, antimicrobial, and virulence profiles of *E. bugandensis*.

## INTRODUCTION

*Enterobacter bugandensis* is a rod-shaped and Gram-negative pathogen that was first isolated from blood samples and powdered milk in a neonatal ward of the Bugando Medical Centre (Mwanza, Tanzania) [1]. Carrying a plasmid-borne *bla*_CTX-M-15_, the lethal EB-247 strain had its genome sequenced, which was the first finished *E. bugandensis* genome (GCF_900324475.1) [2, 3]. Urbaniak et al. [4] later reported the complete genome sequencing of several *E. bugandensis* isolates from the International Space Station (ISS). Notably, all these ISS strains showed resistance to cefazolin, cefoxitin, ciprofloxacin, erythromycin, gentamicin, oxacillin, penicillin, rifampin, and tobramycin, imposing health risks to ISS crew [4].

The massive and often inadequate use of antibiotics have been accelerating the development of antimicrobial resistance (AMR) worldwide [5]. Reports of multidrug-resistant (MDR) *Enterobacter* isolates are on the rise and already reached an emergency level [6], mainly due to the spread of extended-spectrum β-lactamases (ESBLs) and carbapenemases [7].

The increasing availability of genomic data makes *in silico* AMR genotyping a promising strategy in research, surveillance, and clinical decisions [8, 9]. Several authors reported the concordance between genotype and phenotype tests, supporting the use of genomic information in hospitals [10, 11]. In a recent work, Chavda et al. conducted a phylogenomic analysis of *Enterobacter* spp. isolates, revealing the distribution of carbapenem resistance genes (e.g. *bla*_KPC-2_, *bla*_KPC-3_, *bla*_KPC-4_, and *bla*_NDM-*1*_) in the genus [12]. Due to the recent nosocomial outbreaks associated with MDR *Enterobacter* strains [13], new surveillance strategies using high-resolution molecular typing can be employed to determine the population structure and help in epidemiological investigations. Single nucleotide polymorphism (SNP) and gene-by-gene (GBG) approaches have been largely used for bacterial sub-typing [14]. Since SNPs are highly informative markers, their identification in a set of genes conserved across strains (e.g. core genome) is instrumental in revealing evolutionary histories of closely related bacteria [15]. The GBG strategy is based on inference of categorical data of allelic variation from a set of genes [16], commonly the core genome or even the whole genome [17].

In this work, we report the whole-genome sequencing and analysis of a non-clinical *E. bugandensis* strain isolated from cattle manure vermicompost. Through comparative analysis, SNP-based and GBG phylogenetic reconstructions, we reclassified nine published *Enterobacter* spp. genomes as *E. bugandensis.* We analyzed the *E. bugandensis* UENF-21GII genome along with those of publicly available *E. bugandensis* isolates, which allowed us to perform a comprehensive characterization of the genomic content, population structure, virulence, and resistance profiles in this species.

## RESULTS AND DISCUSSION

### Isolation and genome sequencing of a non-clinical *E. bugandensis* strain

During the characterization of culturable bacteria from mature cattle vermicompost, we identified a bacterium that was preliminarily characterized as *Enterobacter* sp. by colony morphology, microscopy, and 16S rRNA sequencing. This isolate was sequenced using an Illumina HiSeq 2500 instrument (paired-end mode, 2 × 100 bp reads). Sequencing reads were processed with Trimmomatic, assembled using SPAdes, and annotated with PGAP (see methods for details). The assembled genome has 17 scaffolds, encompassing 4,765,459 bp, with a GC content of 55.88 %. The genome has 4,539 coding sequences (CDS), 72 tRNA and 11 rRNA genes (Table S1). No plasmids were detected. We used BUSCO to estimate genome completeness and recovered the full set of 781 *Enterobacteriales* single-copy genes, supporting the high quality and completeness of the assembled genome. Average nucleotide identity (ANI) and digital DNA-DNA hybridization revealed that this strain belongs to the *E. bugandensis* species (Figure S1), leading us to name it as *E. bugandensis* UENF-21GII (accession RDOI01000000).

### *E. bugandensis* pangenome determination

Recent works have improved the taxonomic classification of *Enterobacter* spp. [18]. We used the ANI metric to inspect all *Enterobacter* genome assemblies available in Genbank (n = 1616, January 2019) for potentially misclassified genomes (see methods for details). This is an important step that is in line with the recent announcement that NCBI will use a similar strategy in a large scale [19]. This analysis allowed us to reclassify five *E. cloacae* and four *Enterobacter* spp. as *E. bugandensis*, including *E. cloacae* MBRL 1077 (GCA_001562175), originally reported as the first IMI carbapenemase-producing and colistin-resistant *E. cloacae* strain isolated in the USA [20]. Our findings support the recent report from Singh et al. regarding the reclassification of this strain [21].

Next, we conducted a comparative genome analysis of 39 *E. bugandensis* genomes (Table 1). Among the reclassified isolates, two have the same strain name as other *E. bugandensis*: IF2SW-B1* and IF2SW-P2* (GCA_001743375 and GCA_001743385, respectively). We used asterisks to differentiate these isolates from those previously classified as *E. bugandensis* IF2SW-B1 and IF2SW-P2 (GCA_002890755 and GCA_002890725, respectively). Although biased towards clinical and human-associated isolates, non-clinical *E. bugandensis* strains have been described here and elsewhere [22], suggesting that this species occupies a wider range of niches. We used the strains listed above (Table 1) to build the *E. bugandensis* pangenome. Our results support a high genetic diversity and an open pangenome (Figure 1a), features that are often associated with the ability to colonize multiple environments [23]. The *E. bugandensis* pangenome is composed by 11,682 genes families: 3,167 are common to all strains (core genome); 3,785 are present in more than one (but not all) strains (accessory genome) and; 4,730 are unique to any given strain (Figure 1b).

**Figure 1:**
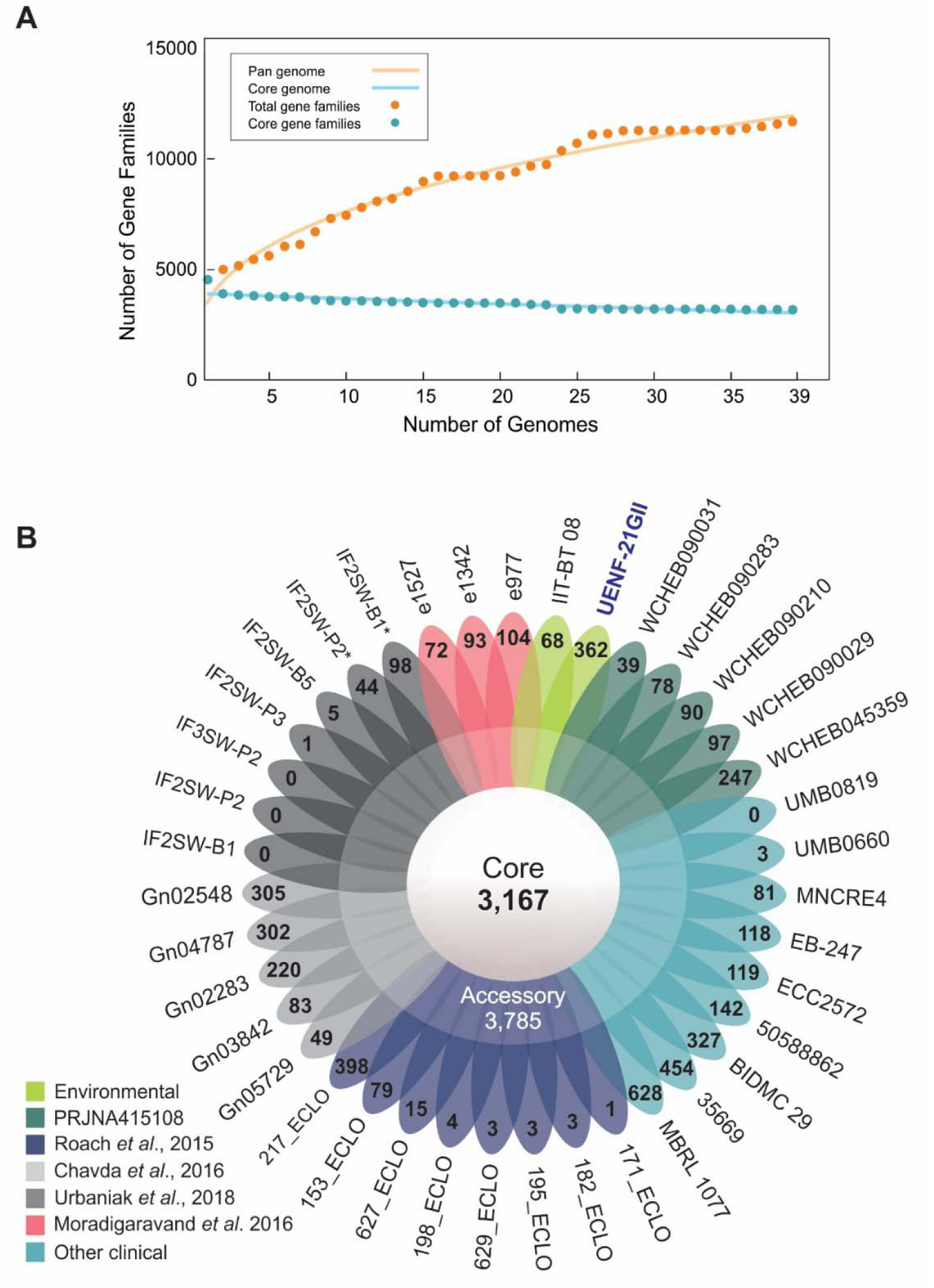
**a)** Number of gene families in the *E. bugandensis* pangenome. The cumulative curve (in orange) supports an open pangenome; **b)** Flowerplot illustrating core and accessory genome sizes, along with the number of unique genes of each *E. bugandensis* strain. Color codes represent original studies or source of the isolates [4, 12, 75, 76].

**Table 1:** *Enterobacter bugandensis* isolates used in this study and their respective ANI against the UENF-21GII strain.

| Accessions | Classified as | ANI (%) | Country | Source |
| --- | --- | --- | --- | --- |
| RDOI01000000_ | *E. bugandensis* UENF-21GII | 100 | Brazil | Vermicompost |
| GCA_003964255 | *E. bugandensis* WCHEB090283 | 98.5 | China | *H. sapiens* |
| GCA_003964645 | *E. bugandensis* WCHEB090031 | 98.5 | China | *H. sapiens* |
| GCA_003964825 | *E. bugandensis* WCHEB090210 | 98.6 | China | *H. sapiens* |
| GCA_003965205 | *E. bugandensis* WCHEB090029 | 98.4 | China | *H. sapiens* |
| GCA_003985275 | *E. bugandensis* WCHEB045359 | 98.6 | China | *H. sapiens* |
| <u>GCA_000515295</u> | *E. cloacae* IIT-BT 08 | 98.5 | India | Plant leaves |
| <u>GCA_000534375</u> | *Enterobacter* sp. BIDMC 29 | 98.0 | Unknown | Unknown |
| GCA_000957065 | *E. bugandensis* 35669 | 98.4 | USA | Unknown |
| GCA_000958885 | *E. bugandensis* MNCRE4 | 98.0 | USA | Urine |
| GCA_001022595 | *E. bugandensis* GN02548 | 99.0 | USA | Bodily fluid* |
| GCA_001023305 | *E. bugandensis* GN02283 | 98.2 | USA | Bodily fluid* |
| GCA_001054435 | *E. bugandensis* 153_ECLO | 98.7 | USA | *H. sapiens* |
| GCA_001054505 | *E. bugandensis* 171_ECLO | 98.2 | USA | Unknown |
| GCA_001054625 | *E. bugandensis* 217_ECLO | 98.6 | USA | Unknown |
| GCA_001055515 | *E. bugandensis* 627_ECLO | 98.9 | USA | *H. sapiens* |
| GCA_001055645 | *E. bugandensis* 182_ECLO | 98.2 | USA | *H. sapiens* |
| GCA_001055805 | *E. bugandensis* 195_ECLO | 98.2 | USA | *H. sapiens* |
| GCA_001055835 | *E. bugandensis* 198_ECLO | 98.2 | USA | *H. sapiens* |
| GCA_001057695 | *E. bugandensis* 629_ECLO | 98.2 | USA | *H. sapiens* |
| <u>GCA_001462935</u> | *Enterobacter* sp. 50588862 | 98.1 | USA | Urine* |
| GCA_001518315 | *E. bugandensis* GN03842 | 97.6 | USA | Blood* |
| GCA_001518395 | *E. bugandensis* GN05729 | 98.2 | USA | Blood * |
| <u>GCA_001562175</u> | *E. cloacae* MBRL 1077 | 95.4 | USA | Skin* |
| GCA_001631115 | *E. bugandensis* GN04787 | 98.5 | USA | Blood* |
| <u>GCA_001743375</u> | *Enterobacter* sp. IF2SW-B1* | 98.5 | - | ISS |
| <u>GCA_001743385</u> | *Enterobacter* sp. IF2SW-P2* | 98.5 | - | ISS |
| GCA_002785865 | *E. bugandensis* ECC2572 | 98.6 | China | Urine |
| GCA_002848105 | *E. bugandensis* UMB0660 | 98.5 | USA | Urine* |
| GCA_002860705 | *E. bugandensis* UMB0819 | 98.5 | USA | Urine* |
| GCA_002890715 | *E. bugandensis* IF3SW-P2 | 98.7 | - | ISS |
| GCA_002890725 | *E. bugandensis* IF2SW-P2 | 98.7 | - | ISS |
| GCA_002890755 | *E. bugandensis* IF2SW-B1 | 98.7 | - | ISS |
| GCA_002890765 | *E. bugandensis* IF2SW-P3 | 98.7 | - | ISS |
| GCA_003627555 | *E. bugandensis* IF2SW-B5 | 98.7 | - | ISS |
| <u>GCA_900075565</u> | *E. cloacae* e1342 | 97.6 | UK | Blood* |
| <u>GCA_900075815</u> | *E. cloacae* e1527 | 98.7 | UK | Blood* |
| <u>GCA_900078165</u> | *E. cloacae* e977 | 98.7 | UK | Blood* |
| <u>GCA_900324475</u> | *E. bugandensis* EB-247 | 98.5 | Tanzania | Blood* |

\* Human samples or tissues. ISS – International Space Station. Reclassified accessions strains are underlined.

*E. bugandensis* UENF-21GII has 362 unique genes, including two genes (EAI35_15025 and EAI35_02745) encoding S-type family pyocin-S domain proteins (a bacteriocin). These toxins induce cell death by DNA cleavage and are commonly associated with niche competition in *Pseudomonas* [24]. Further, we found 27 genomic islands (GIs) in *E. bugandensis* UENF-21GII genome (Table S2), which probably represent recent horizontal gene transfer events. Two GIs (GI1 and GI2) are composed exclusively by unique genes. GI1 has 4,305 bp, encompassing seven genes, mainly of unknown function, while GI2 has 9,148 bp, comprising ten genes related to type 1 fimbrial assembly. Type 1 fimbriae are filamentous surface organelles that mediate bacterial adherence to biotic and abiotic surfaces [25]. Interestingly, *E. bugandensis* UENF-21GII possesses several other genes related to type 1 fimbrial assembly along its core and accessory genomes, suggesting that this unique horizontally-acquired fimbrial operon might play specific roles in this strain. Further, the GI4 comprises curli specific genes (csg) [26] in two divergently transcribed operons (EAI35_03890 to EAI35_03915) downstream of *csgG* (EAI35_03885), which have been linked to host cell adhesion, invasion, and biofilm integrity [27]. Further, the presence of *csg* in the core genome supports curli biofilm assembly as a common virulence strategy in *E. bugandensis*, as reported in other Enterobacteriaceae (reviewed by [28]). Other GIs in *E. bugandensis* UENF-21GII include multiple operons encoding toxin-antitoxin systems and other proteins involved biological conflicts (Table S2). Collectively, these GIs indicate that *E. bugandensis* UENF-21GII is equipped with several horizontally-acquired genes that might be adaptive in highly competitive niches.

We also evaluated the composition of genetic mobile elements in *E. bugandensis* by searching for plasmids and bacteriophage signatures, as these elements can be associated with the emergence of new phenotypic traits. The most frequent bacteriophage was the lambdoid Enterobacteria phage mEp237 (NC_019704), present in 56.41% of the genomes used in this study, and duplicated in four genomes: ECC2572, MBRL1077, WCHEB090029, and WCHEB090283 (Table S3). This high prevalence suggests a role of this bacteriophage in the evolution and diversification of *E. bugandensis*. Regarding the distribution of plasmids, 33.33% of *E. bugandensis* strains had detectable replicon types, including IncX3, IncFII, IncFIA, IncFIB, and Col, being the former the most frequent replicon type (Table S4). Importantly, IncF plasmids are frequently associated with the dissemination of ESBLs, quinolone, and aminoglycoside resistance genes [29].

### Molecular subtyping of *E. bugandensis*

Although serological approaches are a gold standard in bacterial strain typing, such methods sometimes fail in predicting genomic relatedness [30]. Over the past decade, core- and whole-genome multilocus sequencing typing (cgMLST and wgMLST, respectively) have received considerable attention in bacterial epidemiology [17, 31]. We applied both these methods to analyze the population structure of *E. bugandensis*. We used 311,598 SNPs extracted from the 3,167 core genes to perform a maximum-likelihood phylogenetic reconstruction (i.e. cgMLST; Figure 2) to analyze the general population structure of *E. bugandensis*. We recovered five distinct phylogroups (PGs) with high statistical support. Nevertheless, several strains (e.g. *E. bugandensis* UENF-21GII) were not assigned to any PG, indicating that additional groups will be detected when a more diverse collection of *E. bugandensis* genomes become available. An expanded investigation of the *E. bugandensis* population structure using GBG wg and cgMLST dendrograms (Figure S2 and S3, respectively) also supported the clustering patterns observed above, with the exception of the PG-C group. The slight incongruences between GBG- and SNP-based approaches are not surprising given the nature and assumptions of these methods. The long branch length of MBRL 1077 in the SNP cgMLST phylogenetic tree (Figure 2) supports this strain as a representative isolate of PG-C, which was poorly represented among publicly available genomes.

**Figure 2:**
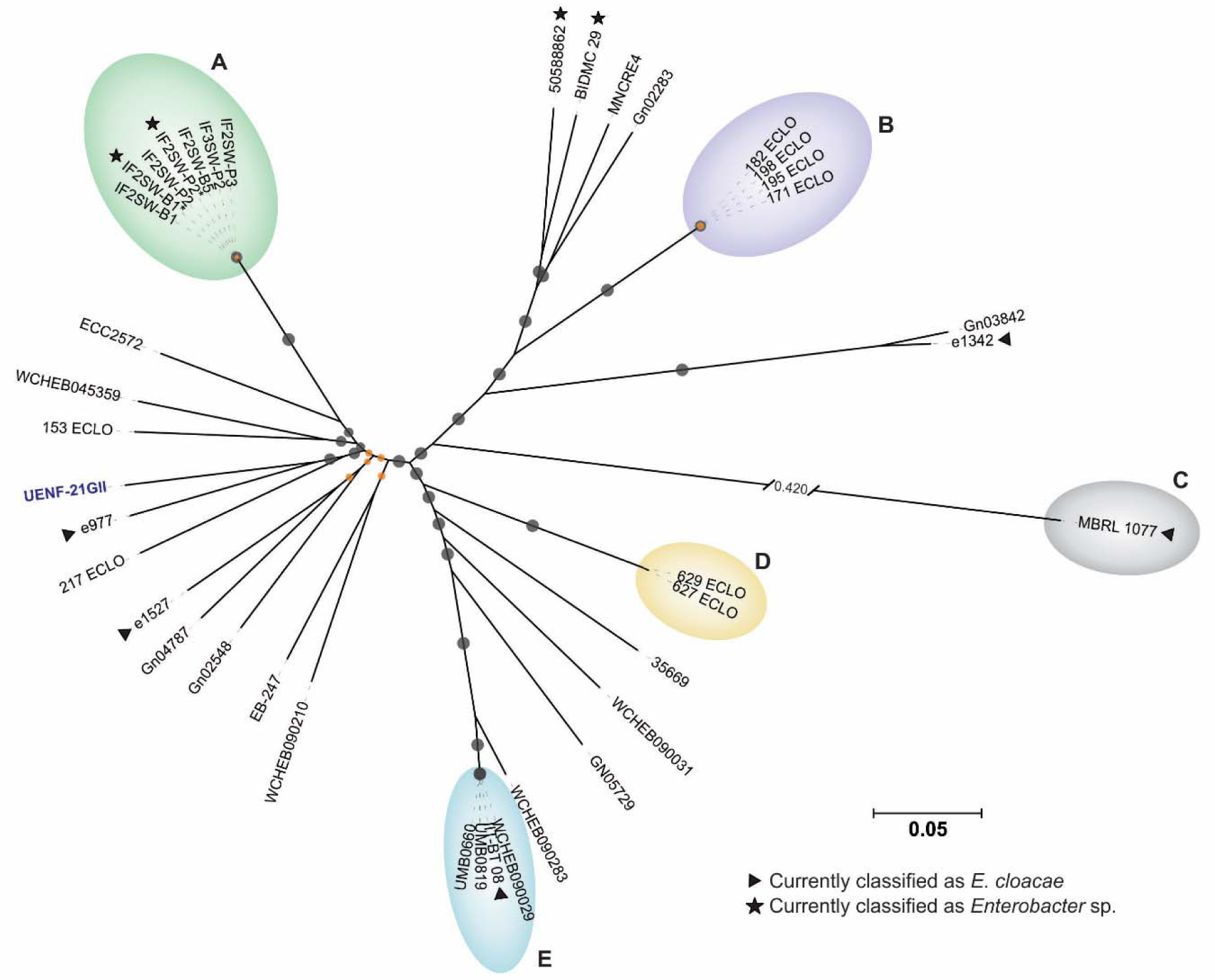
Molecular subtyping of *E. bugandensis* showing phylogenetic groups (PG-A to PG-E). SNPs extracted from the core-genome were used to build a maximum likelihood phylogenetic tree using RAxML (see methods for details). Bootstrap values below and above 60 % are represented by orange and gray circles, respectively. Arrowheads and stars represent reclassified *E. cloacae* and *Enterobacter* spp. strains, respectively.

PG-A is the largest PG and comprises only isolates from ISS surveillance initiative [4], supporting a common origin of these strains. The PG-E group comprises the environmental isolate from India (II-BT 08), along with clinical isolates from China (prefix WCHEB) and U.S. (prefix UMB). This heterogeneous group highlights the worldwide dissemination and diverse lifestyle of *E. bugandensis*. The admixture between environmental and clinical samples will be better understood when more genomes are sequenced. In a recent work, Singh et al. [21] stated, based on whole genome alignments, that ISS isolates formed an unique ecotype along with the type strain EB-247, 153_ECLO and MRBL 1077. Here, we performed a broader phylogenetic reconstruction of *E. bugandensis*, revealing that ISS and clinical isolates do not belong to the same PG. In addition, our analysis confirms the outgroup behavior of MBRL 1077 strain.

### Distribution of antimicrobial resistance and virulence genes

The distribution of AMR and virulence genes across *E. bugandensis* strains remain poorly understood. We have further analyzed the *E. bugandensis* core genome to identify the core resistome and core virulome. A total of 96 and 52 core resistance and virulence genes were found, respectively (Table S5 and S6). We mapped these genes using the *E. bugandensis* UENF-21GII genome as reference (Figure 3). The presence of the enterobactin operon *entABCDEFS*, flanked by *fep* genes, implies that iron uptake via siderophore production [32] is essential in *E. bugandensis*. Also related to iron acquisition, the operon *iucABCD*/*iutA* is present in the *E. bugandensis* core virulome. This operon encodes the aerobactin system, which plays key roles during intracellular growth [33]. Pati et al. (2018) identified 141 up-regulated genes in *E. bugandensis* EB-247 (GCF_900324475.1) cultured in the presence of human blood serum [2]. This gene set includes genes encoding different iron acquisition systems, such as enterobactin (*ent*) and aerobactin (*iuc*) [2]. The presence of these genes in the core virulome demonstrates the importance of iron chelation in the virulence of *E. bugandensis*.

**Figure 3:**
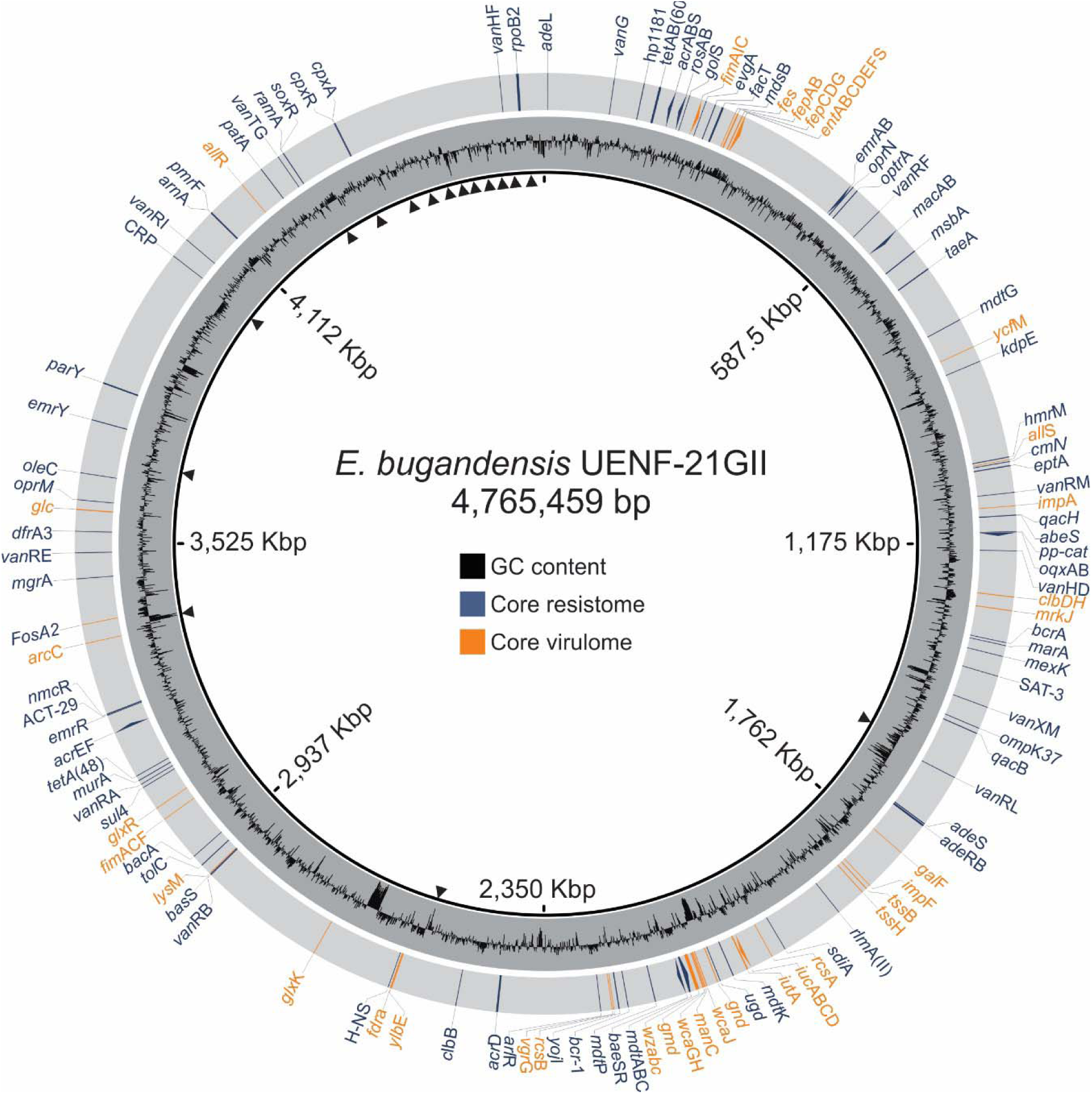
Core virulome and resistome genes mapped in the *E. bugandensis* UENF-21GII genome. Arrowheads represent scaffold ends. Blue and orange genes represent core resistome (95 genes) and virulome (52 genes). The core resistome includes genes conferring resistance to cephamycin, cephalosporin, penam, and carbapenems. Many efflux pumps are also present (e.g. *oqx*AB, *ros*AB, *acr*AB-TolC). The core virulome harbors the siderophore production operons *ent*ABCDEFS (enterobactin) and *iuc*/*iut* (aerobactin), which are marked with asterisks.

The core resistome is composed of a large number of efflux pumps (e.g. OqxAB, RosAB, AcrAB-TolC), which seems to be the main intrinsic resistance mechanism of *E. bugandensis* (Table S5). Further, we found that ACT-29 is the chromosomal constitutive intrinsic AmpC β-lactamase in *E. bugandensis*; ACT-29 is typically associated with resistance against cephamycin, cephalosporin, penam, and carbapenem [34]. Other core genes were predicted to confer resistance against aminocoumarin, aminoglycosides, fosfomycin, fluoroquinolones, and macrolides (Table S5). In addition to genes that are directly involved with AMR, we also identified global metabolic regulators (e.g. H-NS, BaeRS, and marA), which modulate the transcription of an array of AMR genes [35, 36]. In Enterobacteriaceae, the histone-like protein H-NS binds to AT-rich promoter regions, preferentially silencing horizontally-transferred genes [37]. H-NS has been shown to modulate virulence [38], motility [39], capsule biosynthesis [40], and expression of MDR efflux pumps [41].

Next, we screened the content of accessory and unique genomes. To illustrate the gene presence/absence profile of each strain, we mapped the resistance genes on the SNP-cgMLST tree (Figure 4). Many acquired resistance genes potentially confer resistance to quinolones (*qnr*B17), aminoglycosides (*aadA*), fosfomycin (*fosA*), and macrolides (*srmB*) in *E. bugandensis* (Table S7). The *bla*_IMI-1_ carbapenemase gene was found in two strains (50588862 and MBRL 1077). In addition, we report for the first time the presence of the New Delhi metallo-β-lactamase-5 (*bla*_NDM-5_) co-occurring with *bla*_CTX-M-55_ in *E. bugandensis* (strain WCHEB090029; Figure 4, Table S4).

**Figure 4:**
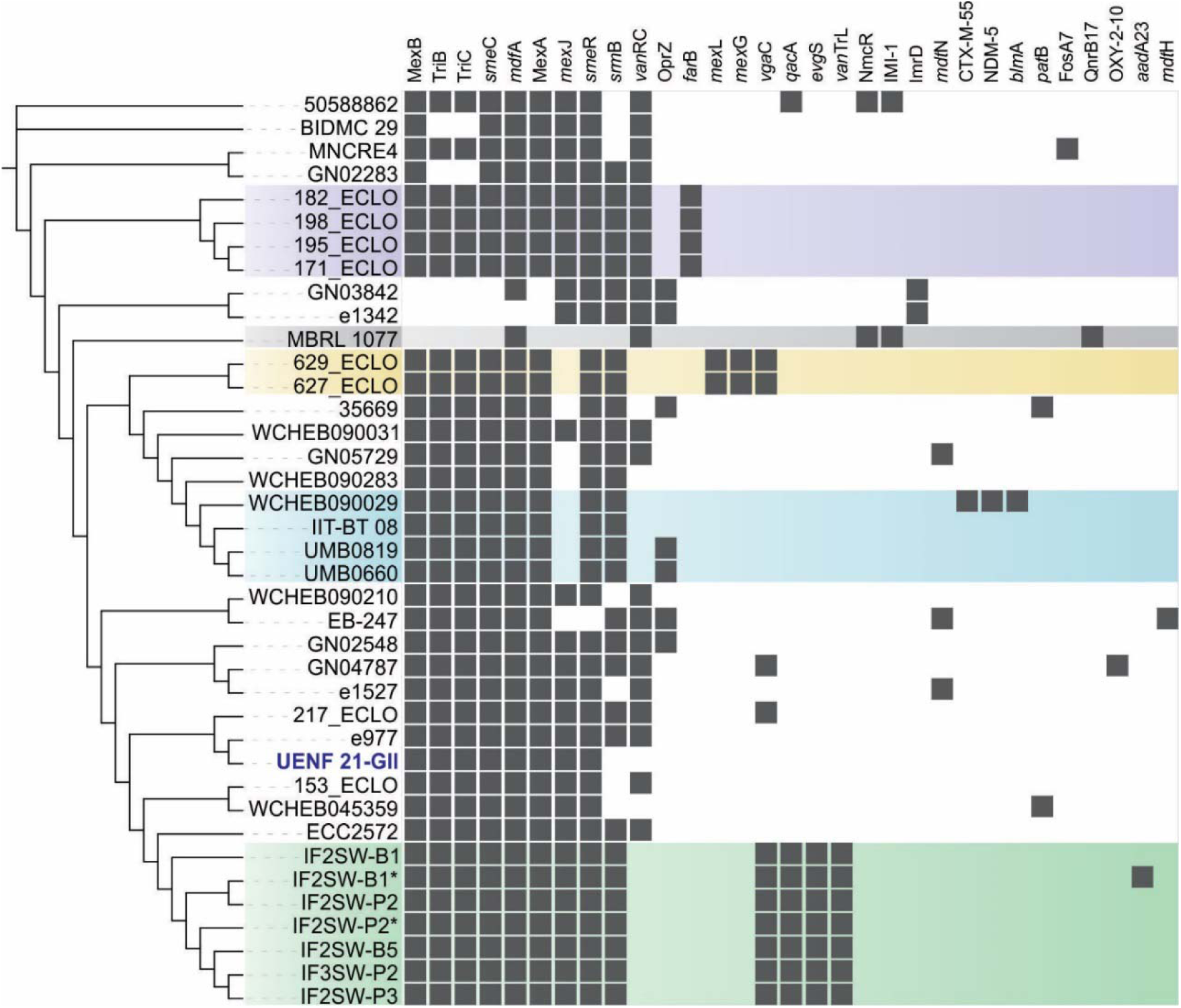
Accessory resistome of *E. bugandensis*. Highlights represent the phylogroups found in the population structure analysis (see Figure 2). There are four genes (i.e. *vgaC, qacA, evgS* and *vanTrL*) shared by (but not exclusive of) all ISS strains (PG-A, green). We also found two genes (*mexL* and *mexG*) related to an efflux pump in PG-D (yellow). The ESBLs *bla*_IMI-1_ (in MRBL 1077 and 50588862 strains) and *bla*_CTX-M-55_ and *bla*_NDM-5_ (in WCHEB090029) are also of notice. Although extensively documented elsewhere [2], the sequence of the *E. bugandensis* EB-247 plasmid (pEB-247, accession number LN830952) is not linked to the EB-247 genome accession (GCA_900324475). Although not included in our analysis, this plasmid carries importance acquired resistance genes (e.g. *bla*_CTX-M-15_, *bla*_TEM-1_, *bla*_OXA-1_).

The *bla*_NDM-5_ gene encodes one of the most clinically-relevant carbapenemases and has been previously reported within IncX plasmids in *Escherichia coli* [42]. The protein encoded by *bla*_NDM-5_differs by only two amino acids from that encoded by *bla*_NDM-1_ (Val88Leu and Met154Leu) [43] and confers a broad-spectrum hydrolytic activity against penicillins, cephalosporins and carbapenems (except monobactams) [44]. In an elegant study, Ali et al. (2018) showed that the hydrolytic activity of NDM-5 is stronger than that of NDM-1, −4, −6 and −7, exhibiting a four-fold increase in the minimum inhibitory concentration of imipenem and meropenem when compared to NDM-1 [45]. Further, through molecular docking, Ali et al. proposed that NDM-5 is the only variant capable of creating a stable interaction with imipenem and meropenem. Plasmids from a variety of incompatibility groups have been reported to carry *bla*_NDM-5_, typically along with several other AMR genes [46]. IncX3 plasmids were involved with *bla*_NDM_ dispersion in *Enterobacteriaceae* in China [47] and South Korea [48].

Genetic context analysis of *bla*_NDM-5_ in the IncX3 plasmid of WCHEB090029 showed that this gene belongs to a 11,815 bp operon (*umu*D-IS26-*dsb*D-*trp*F-*ble*-*bla*_NDM-5_-IS5-ISAba125-IS3000) that is flanked by two transposases (ISKox3 and Tn2). This operon exhibits the same architecture found in four other IncX3 plasmids from other Enterobacteriaceae strains isolated in China [49]. The identification of this plasmid in *E. bugandensis*, together with the presence of this species in several niches, indicates that the spread of plasmid-associated *bla*_NDM-5_ to other species constitutes a real risk that warrants further surveillance.

The *E. bugandensis* accessory virulome (Table S8) shows that the PG-A, composed only by ISS isolates (Figure 2), possesses extra capsule production genes (i.e. *kvg*AS) (Figure 5). Interestingly, we also noticed the presence of the iron transport operon *iro*BCDEN across eleven (28.2 %) strains. Originally identified in *Salmonella*, this C-glucosylated form of enterobactin was termed salmochelin, and is often found in uropathogenic *E. coli* (UPEC) [50]. Salmochelins are more efficient than enterobactins in promoting bacterial growth [51], suggesting that they might confer a competitive advantage to *iro*-harboring *E. bugandensis* strains in particular conditions. Notably, all PG-E isolates harbor the full *iro*BCDEN operon (Figure 2), indicating that this region might have been horizontally transferred to the common ancestor of this group. Thus, we investigated the presence of GIs in the only complete genome available in the PG-E (II BT-08; accession GCA_000515295 [22]). GI3, encompassing 11,148 bp, harbors the entire *iro*BCDEN operon, confirming the horizontal transfer of this region. We also found other 31 putative GIs in this strain (Table S9).

**Figure 5:**
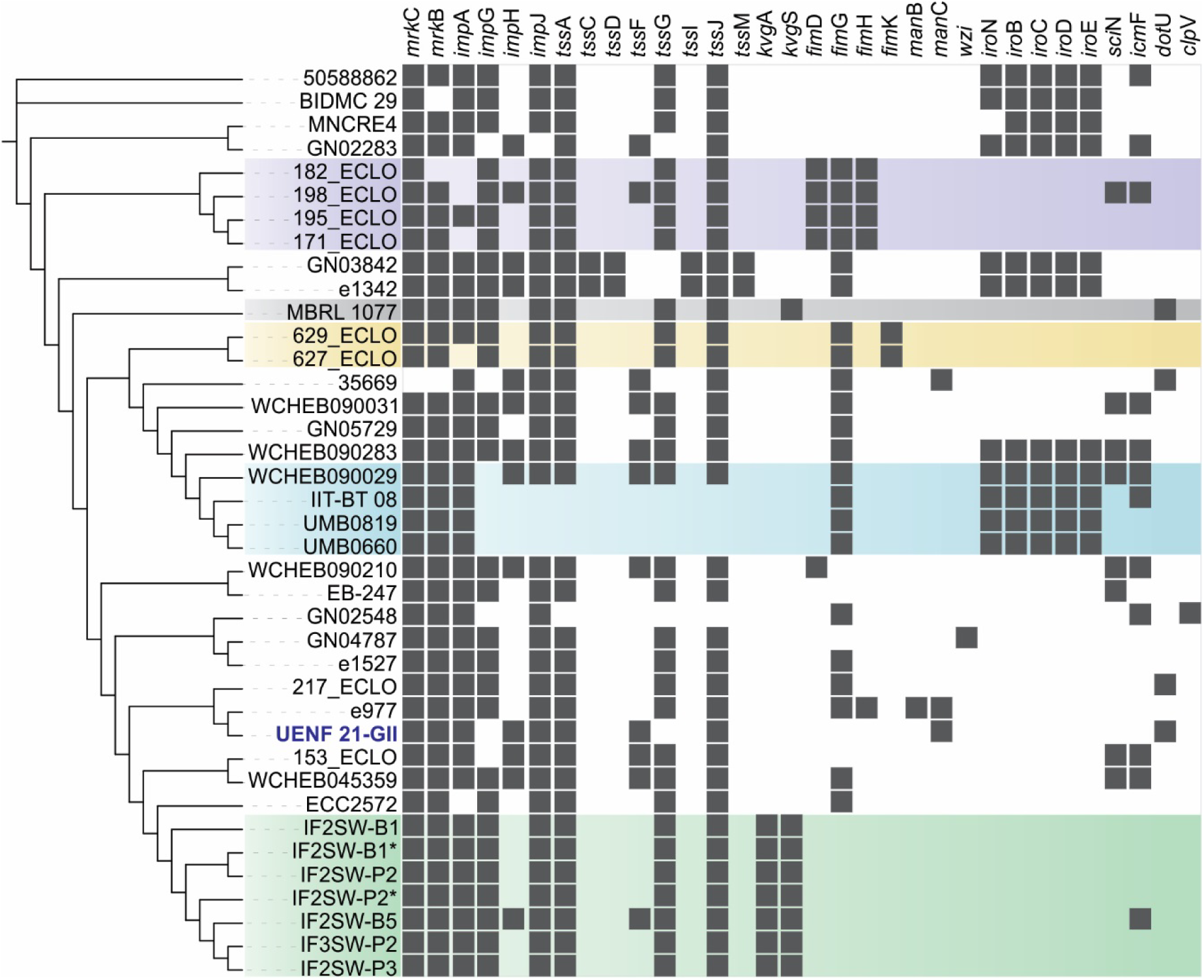
Accessory virulome of *E. bugandensis*. Highlights represent the phylogroups found in the population structure analysis (see Figure 2). The main findings include the presence of an additional capsule system (*kvg*AS) in ISS isolates (PG-A, in green) and the presence of a salmochelin (*iro*BCDEN) operon across 11 strains, including all PG-E isolates (in blue).

Our results reveal an alarming antibiotic resistance profile in *E. bugandensis*, especially due to the presence of ESBLs. We also uncovered a sophisticated iron acquisition system and showed the presence of a salmochelin operon that is conserved in several strains, including all PG-E isolates, further supporting the monophyly of this PG. In addition, we showed that the ISS isolates belong to a separate phylogroup (PG-A) and share genes involved in capsule production. We also report, for the first time in *E. bugandensis*, the co-occurrence of the ESBLs *bla*_CTX-M-55_ and *bla*_NDM-5_. Collectively, this work provides a genomic foundation for further developments towards epidemiological surveillance and biological understanding of this emerging pathogen that can colonize several niches and potentially spread AMR resistance genes.

## METHODS

### Methods vermicompost maturation

Mature vermicompost was produced inside a 150 L cement ring using dry cattle manure as substrate. Humidity was kept at 60-70%, by weekly watering and mixing. After 1 month, earthworms (*Eisenia foetida*) were introduced at the rate of 5 kg·m^3^. After 4 months, earthworms were removed and the stable vermicompost was stored in plastic bags at 25 °C. At the final maturation stage, the chemical composition of the substrate (in g·kg^−1^) was as follows: total nitrogen (1.9 ± 0.4); total carbon (22.99 ± 3.3); P_2_O_5_ (6.97 ± 1.4); C/N ratio of 13.8 ± 0.4 and pH (H_2_O) = 6.6 ± 0.18.

### Bacterial isolation and DNA purification

Serial dilutions were performed on a solution prepared by the addition 10 g of vermicompost in 90 mL of saline (8.5 g·L^−1^ NaCl), followed by shaking for 60 min. Next, 1 mL of the initial dilution (10^−1^) was added to a new tube containing 9 mL of saline (10^−2^). This procedure was repeated until reaching a 10^−7^ dilution. Then, 100 μL of the final dilutions from 10^−5^ to 10^−7^ were spread on plates containing solid Nutrient Broth (NB) with 8 g·L^−1^ of NB and 15 g·L^−1^ of agar in 1 L of distilled water. After incubation at 30 °C for 7 days, individual colonies were transferred for purification to Petri plates with Dygs solid media [52] containing 2 g·L^−1^ of glucose, 2 g·L^−1^ of malic acid, 1.5 g·L^−1^ of bacteriological peptone, 2 g·L^−1^ of yeast extract, 0.5 g·L^−1^ of K_2_HPO_4_, 0.5 g·L^−1^ of MgSO_4_·7H2O, 1.5 g·L^−1^ of glutamic acid and 15 g·L^−1^ of agar, adjusted to pH 6.0; these supplies were acquired from Vetec (São Paulo, Brazil). From the last dilution (10^−7^) and after the isolation and purification on Dygs solid medium, a brown, circular, punctiform and smooth surface bacterial colony was selected. Phase contrast microscopy revealed the presence of rod-shaped motile cells and Gram-negative stain under light microscopy. This distinctive isolate, named UENF-21GII, was stored in a 16 mL glass flask containing 5 mL of Nutrient Broth solid medium covered with mineral oil and, later, grown in liquid Dygs medium using a rotatory shaker at 150 rpm and 30 °C for 36 h. Total DNA of the UENF-21GII isolate was extracted using QIAamp® DNA Mini Kit (QIAGEN GmbH, Hilden, Germany). DNA quantification and quality assessment were performed using an Agilent Bioanalyzer 2100 instrument (Agilent, California, USA).

### Genome sequencing and assembly

The sequencing procedures and basic data processing were performed as previously described [53]. In summary, paired-end libraries were prepared with the TruSeq Nano DNA LT Library Prep (Illumina) and sequenced on a HiSeq 2500 instrument at the Life Sciences Core Facility (LaCTAD; UNICAMP, Campinas, Brazil). The quality of the sequencing reads (2 × 100 bp) was checked with FastQC 0.11.5 (https://www.bioinformatics.babra-ham.ac.uk/projects/fastqc/). Quality filtering was performed with Trimmomatic 0.35 [54] and reads with average quality below 30 were discarded. The *E. bugandensis* UENF-21GII genome was assembled with SPAdes 3.12.0 [55] and scaffolded with Gfinisher 1.4 [56], using as references the complete genome of *E. bugandensis* EB-247 (GCA_900324475.1) and an alternative assembly generated with Velvet 1.2.10 [57]. QUAST 5.0.1 [58] was used to assess general assembly statistics. Genome completeness was inspected with BUSCO 3.0 [59], using the *Enterobacteriales* dataset as reference.

### Genome annotation and phylogenetic analysis

The assembled genome was annotated with the NCBI Prokaryotic Genome Annotation Pipeline [60]. The presence of plasmids was assessed with plasmidSPAdes 3.12 [61] and PlasmidFinder 1.3 [62]. Bacteriophage signatures were analyzed with PHASTER [63]. Genes involved in antibiotic resistance and virulence were predicted by BLAST [64] searches against the CARD database 2.03 [65] and the VFDB database [66] complemented with other reference virulence genes curated from the literature [67]. For BLAST searches, we used 50 % and 80 % identity and coverage thresholds, respectively. The *E. bugandensis* UENF-21GII genome was deposited on Genbank (BioProject: PRJNA497272, accession RDOI00000000). Publicly available genomes were downloaded and protein sequences were predicted using PROKKA 1.13.3 [68]. Pangenome analysis was performed with BPGA 1.3.0, applying an identity cutoff of 95 % [69]. Protein sequences were aligned MUSCLE 3.8.31 (default parameters) [70]. SNPs were extracted using SNP-sites 2.3.3 [71] and maximum-likelihood phylogenetic reconstructions performed using RaxML 8.2.12 [72], using the ASC_GTRGAMMA, Paul Lewis standard correction and 1000 bootstrap replicates. PGAdb was used to build cg- and wgMLST by GBG, generating 3,548 and 11,170 loci respectively [31]. Phylogenetic trees and gene presence/absence profiles were integrated and rendered using iTOL 3 [73]. Pyani 0.27 was used to calculate ANI between all 1,616 genome assemblies deposited under the *Enterobacter* genus (as of January of 2019). GIs were predicted using IslandViewer 4 [74], followed by manual inspection.

## ACKNOWLEDGEMENTS

This work was supported by Fundação Carlos Chagas Filho de Amparo à Pesquisa do Estado do Rio de Janeiro (FAPERJ; grant E-26/102.259/2013) and by Conselho Nacional de Desenvolvimento Científico e Tecnológico (CNPq; grant 449904/2014-8). FPM and FP-S postgraduate fellowships were funded by Coordenação de Aperfeiçoamento de Pessoal de Nível Superior (CAPES; Finance Code 001). FPM postdoctoral fellowship and HP-A undergraduate fellowship were funded by UENF. We would like to thank the staff of the Life Sciences Core Facility (LaCTAD) of UNICAMP for library preparation and genome sequencing.

