## Supplementary material for "Population structure and pangenome analysis of Enterobacter bugandensis uncover the presence of *bla*_CTX-M-55_, *bla*_NDM-5_ and *bla*_IMI-1_, along with sophisticated iron acquisition strategies": Figure S1

| | *E. bugandensis* UENF-21GII | *E. bugandensis* EB-247 | *E. asburiae* ATCC 35953 | *E. cancerogenus* ATCC 33241 | *E. chengduensis* WHECI-C4 | *E. cloacae* ATCC 13047 | *E. hormaechei* ATCC 49162 | *E. kobei* DSM 13645 | *E. kobei* JCM 8580 | *E. ludwigii* EN-119 | *E. mori* LMG 25706 | *E. muelleri* JM-458 | *E. soli* ATCC BAA-2102 | *E. timonensis* mt20 |
|---|---|---|---|---|---|---|---|---|---|---|---|---|---|---|
| *E. bugandensis* UENF-21GII | | 98.5 | 91.2 | 86 | 90.3 | 87.7 | 87.7 | 90.4 | 78.7 | 87.6 | 88.5 | 91.1 | 85.3 | 82.1 |
| *E. bugandensis* EB-247 | 89.3 | | 91.3 | 86.1 | 90.3 | 87.8 | 87.7 | 90.5 | 79 | 87.6 | 88.4 | 91.1 | 85.5 | 82.1 |
| *E. asburiae* ATCC 35953 | 45 | 45.1 | | 86.4 | 92.8 | 88.5 | 87.5 | 90.5 | 79.3 | 88.3 | 89.6 | 97.1 | 85.9 | 82.5 |
| *E. cancerogenus* ATCC 33241 | 31.7 | 31.8 | 32.4 | | 85.6 | 85.3 | 85.6 | 85.5 | 78.7 | 85.1 | 85.8 | 86 | 84.7 | 81.7 |
| *E. chengduensis* WHECI-C4 | 42.6 | 42.4 | 52.4 | 31.9 | | 87.6 | 86.8 | 90.3 | 79.1 | 87.4 | 88.3 | 92.6 | 85.2 | 82 |
| *E. cloacae* ATCC 13047 | 35.4 | 35.6 | 37.2 | 30.8 | 35.9 | | 86.3 | 87.6 | 79 | 87.2 | 87.1 | 87.8 | 85.1 | 81.6 |
| *E. hormaechei* ATCC 49162 | 35.5 | 35.7 | 34.6 | 31.6 | 34.2 | 32.9 | | 87 | 79.3 | 85.9 | 87.2 | 87.1 | 85.3 | 82.3 |
| *E. kobei* DSM 13645 | 42.8 | 43.1 | 42.7 | 31.2 | 43 | 35.7 | 33.7 | | 79.3 | 87.5 | 87.8 | 90.2 | 85.1 | 81.9 |
| *E. kobei* JCM 8580 | 22.8 | 23.2 | 23.3 | 22.7 | 23.1 | 23.3 | 23.2 | 23.4 | | 78.6 | 78.6 | 79 | 78.5 | 78.8 |
| *E. ludwigii* EN-119 | 34.5 | 34.8 | 36.1 | 30.4 | 34.7 | 34.2 | 31.6 | 34.6 | 22.6 | | 86.7 | 88 | 85.2 | 81.3 |
| *E. mori* LMG 25706 | 37 | 37 | 39.3 | 31.9 | 37.2 | 34 | 34.2 | 35.4 | 22.6 | 32.6 | | 89 | 85.4 | 81.4 |
| *E. muelleri* JM-458 | 44.4 | 44.7 | 77 | 32.2 | 51.8 | 36.3 | 34.2 | 41.8 | 23 | 35.5 | 39.3 | | 85.7 | 89 |
| *E. soli* ATCC BAA-2102 | 30.5 | 30.5 | 31.2 | 29.4 | 30.6 | 30.3 | 30.2 | 30.1 | 22.2 | 30.3 | 30.8 | 31 | | 81.1 |
| *E. timonensis* mt20 | 26 | 26 | 26.3 | 25.5 | 26 | 25.6 | 36.1 | 25.5 | 22.6 | 25.1 | 25.5 | 26 | 24.7 | |

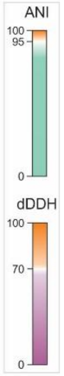


**Figure S1:** Genome relatedness of *E. bugandensis* UENF-21GII and type strains of the *Enterobacter* genus. Average nucleotide identity (ANI) and digital DNA-DNA hybridization (dDDH) values are shown in green and purple, respectively. Values above the species minimal cutoffs (i.e. ANI ≥ 95 and dDDH ≥ 70) are represented in orange.
