## Supplementary material for "Population structure and pangenome analysis of Enterobacter bugandensis uncover the presence of *bla*_CTX-M-55_, *bla*_NDM-5_ and *bla*_IMI-1_, along with sophisticated iron acquisition strategies": Figure S2

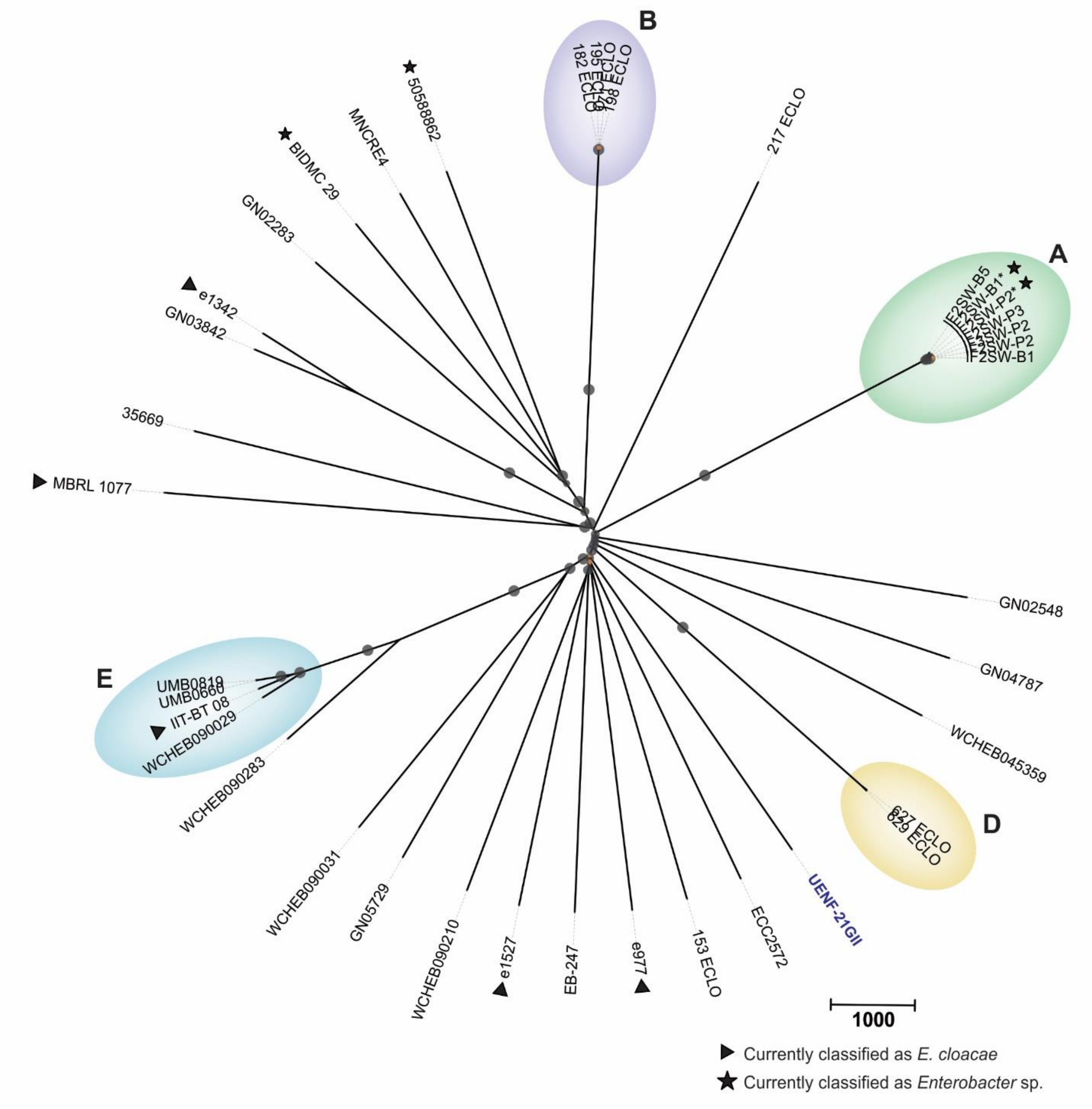


**Figure S2:** Whole genome multilocus sequence typing (wgMLST) dendrogram built with UPGMA in PGAdb. Bootstrap values below 60 % are represented by orange circles; gray circles represent bootstrap values greater than 60 %. Arrowheads and stars depict reclassified strains from *E. cloacae* and *Enterobacter* spp. respectively
